# Chronic infection control relies on T cells with lower foreign antigen binding strength generated by N-nucleotide diversity

**DOI:** 10.1101/2022.06.26.497644

**Authors:** Hassan Jamaleddine, Dakota Rogers, Geneviève Perreault, Judith N. Mandl, Anmar Khadra

**Author notes:** Corresponding authors: Judith Mandl and Anmar Khadra, **Email:** and. These authors contributed equally (Hassan Jamaleddine, Dakota Rogers). These authors contributed equally (Judith N. Mandl, Anmar Khadra).

## Abstract

The pathogens to which T cells respond is determined by the T cell receptors (TCRs) present in an individual’s repertoire. Although more than 90% of the TCR repertoire is generated by terminal deoxynucleotidyl transferase (TdT)-mediated N-nucleotide addition during V(D)J recombination, the benefit of TdT-modified TCRs remains unclear. Here, we computationally and experimentally investigated whether TdT systematically modifies the affinity distribution of a TCR repertoire in ways that impacts acute or chronic infection. Our computational model predicts a shift toward low-affinity T cells over time during chronic, but not acute, infections. Elimination of low-affinity T cells *in silico* substantially delayed chronic infection clearance. Corroborating an affinity-centric benefit for TCR diversity, we showed that infection of TdT-deficient mice delayed the clearance of a chronic viral pathogen, while acute viral control was unaffected. Our data thus suggest that TdT-mediated TCR diversity is of particular benefit in the control of prolonged pathogen replication.

## Introduction

The generation of lymphocyte receptor diversity is a key feature of adaptive immunity (Cooper and Alder, 2006; Schatz and Ji, 2011). For T cells, this diversity is established by somatic recombination of the V(D)J gene segments that constitute the α- and β-chains of the T cell receptor (TCR) (Schatz and Ji, 2011). The T cell response to any one pathogen consists of a number of T cell clonotypes, each expanded from a rare antigen-specific T cell defined by the unique TCR they express. Every T cell clonotype in a given anti-pathogen response recognizes the same or different antigens in the form of peptides presented by major histocompatibility complexes (pMHC) (Yassai et al., 2009) and T cell clonotypes can differ in their ligand binding strengths by several orders of magnitude (Andargachew et al., 2018; Kolawole et al., 2020). Recent evidence suggests that heterogeneity among responding T cells in their TCR binding affinity to pMHC, henceforth referred to as pMHC reactivity, correlates with important differences in effector function. For instance, pMHC binding strength in CD4^+^ T cells has been shown to impact early effector lineage differentiation (DiToro et al., 2018; Snook et al., 2018; Van Panhuys, 2016), while among CD8^+^ T cells it correlates with both their ability to induce target cell lysis and proliferative capacity, and also impacts memory T cell development (Kolawole *et al*., 2020). However, to what extent the pMHC-reactivity distribution of the T cells that constitute a given response affects how quickly a pathogen can be cleared remains incompletely understood.

Experimental techniques are currently limited in their ability to comprehensively study the temporal evolution of T cell clonotype frequencies with distinct pMHC reactivities that make up the antigen-specific response. One common method of identifying antigen-specific T cells is by pMHC-tetramers, which primarily tag clonotypes on the higher end of the pMHC-reactivity spectrum, while missing most low affinity T cells (Andargachew *et al*., 2018). Employing pMHC-tetramers also requires *a priori* knowledge of the epitope recognized by the T cell population being investigated – tracking only the T cells that are specific to one epitope rather than the entire population of responding T cells. Two-dimensional (2D) binding assays measuring TCR-ligand binding affinity similarly rely on knowing the specific epitope recognized by the T cells under study (Huang et al., 2010). Tracking T cell responses by focusing on only a subset of epitope-specific T cells can therefore introduce biases and disregards the contribution of the remaining pathogen-specific T cell population. Complementing experimental results with a theoretical framework that accounts for pMHC reactivity in a T cell repertoire is thus a useful approach to obtaining a clearer understanding of the mechanisms impacting pMHC-reactivity profiles and, consequently, T cell responses to infection.

A critical contribution to the diversification of the T cell repertoire in all jawed vertebrates is made by a DNA polymerase, called terminal deoxynucleotidyl transferase (TdT), that adds non-templated nucleotides to the V(D)J junctions in αβTCRs (Cabaniols et al., 2001; Gilfillan et al., 1995b; Litman et al., 2010; Schatz and Ji, 2011), enhancing TCR repertoire diversity ∼10 fold from the germline recombinatorial diversity alone (Davis and Bjorkman, 1988; Murugan et al., 2012; Zarnitsyna et al., 2013). However, the benefit of the N-diversification mediated by TdT has remained elusive given that TdT-knockout (KO) mice have shown no increased susceptibility to infection, nor any detectable impairment in their response to challenge with an acute pathogen (Gilfillan et al., 1995a; Gilfillan *et al*., 1995b). Interestingly, the genetic sequence and structure of TdT is highly conserved across vertebrates (Hansen, 1997; Lee and Hsu, 1994), suggesting a hitherto unclear evolutionary benefit for its mechanism of action. One hypothesis proposed is that TdT introduces TCRs that, on average, possess lower reactivity to foreign pMHC (Vrisekoop et al., 2014). Indeed, it was shown with influenza A virus infection that the HA_518_ epitope-specific CD8^+^ splenic T cells from TdT KO mice were about 10 times more sensitive to antigenic stimulation as measured by IFNγ production than epitope-specific CD8^+^ T cells from wild type mice, in line with the idea that TdT-independent TCRs may have higher ligand affinity (Haeryfar et al., 2008).

Moreover, during chronic antigen stimulation in infection and cancer mouse models alike, T cells with higher pMHC-reactivity have been shown to be more prone to exhaustion, whereby their cytokine production and contribution to pathogen or tumour control is substantially impaired over their low-affinity counterparts (Alexander-Miller et al., 1996; Shakiba et al., 2021; Wherry et al., 2003). Thus, TdT-dependent TCRs may confer an advantage during chronic infections if the more germline TCRs with higher pMHC-reactivity are more likely to become exhausted and ineffective.

Here, we sought to generate a theoretical framework to examine the role of heterogeneity in T cell reactivity to foreign pMHC in the clearance of an acute or chronic pathogen. To predict the impact of a TdT-deficient T cell repertoire on infection outcomes, we developed and implemented a computational model capturing the kinetics of both acute and chronic pathogen replication that also explicitly considered the evolution of TCR affinity distributions during infection. Our simulations showed that, during chronic infection, there was a decrease in the average pMHC reactivity of the antigen-specific T cell population. Importantly, our model suggested that when the TCR repertoire lacked T cells with lower pMHC reactivity, pathogen clearance during chronic, but not acute, infection was delayed. In line with these model predictions, chronic lymphocytic choriomeningitis virus (LCMV) infection of mice with TdT KO T cells led to more protracted viral replication and a significant delay in viral clearance. Taken together, our modeling and experimental results support the notion that one evolutionary benefit of TdT may be to increase the frequency of T cells with lower pMHC reactivity, and in doing so providing better control of ongoing pathogen replication during chronic infection and preventing them from persisting for longer periods of time within the host.

## Results

### T cell population model captures kinetics of viral load for both acute and chronic infection

In order to compare the temporal evolution of pMHC reactivities of responding T cell clonotypes during acute and chronic infections, we first needed a simple computational model that could recapitulate the kinetics of both rapidly cleared and prolonged pathogen replication. We developed a model that could replicate the serum viral loads obtained upon infection with two strains of LCMV that differ by only 3 coding point mutations, Armstrong (LCMV-Arm) and Clone 13 (LCMV-Cl13), and produce the time course of acute and chronic viral loads in the host, respectively (Ahmed et al., 1984; Bergthaler et al., 2010; Wherry *et al*., 2003). Importantly, the single amino acid mutations between LCMV-Arm and -Cl13 do not occur in peptide segments from which known T cell specific pMHC epitopes are derived (Abdel-Hakeem, 2019; Kotturi et al., 2007) and thus different infection outcomes are governed exclusively by viral replication dynamics within infected cells (Abdel-Hakeem, 2019; Bergthaler *et al*., 2010). Our model considered key interactions between the pathogen load (*P*) and the pMHC-reactivity continuum of all responding effector CD4^+^ and CD8^+^ T cells (*E*) (**Figure 1A**). The dynamics of *P* and *E* depend on the rates of pathogen replication, T cell proliferation and expansion upon antigen encounter, thymic input, homeostatic T cell turnover, and pathogen-induced exhaustion and/or cell death (**Figure 1A**). To incorporate pMHC reactivity explicitly into our model, we defined the thymic input into the effector T cell pool, σ_*E*_, to be a function of pMHC reactivity (denoted *a*_*k*_, **Figure 1B**), a parameter proportional to the strength of TCR signaling (refer to Methods) whereby *a*_*k*_ represents how likely it is that a T cell will proliferate upon antigenic stimulation (Standifer et al., 2009). Consistent with previous studies, we assumed that the strength of negative regulatory mechanisms (such as T cell exhaustion and activation-induced cell death) is positively correlated with the pMHC reactivity of a given T cell (Alexander-Miller *et al*., 1996; Shakiba *et al*., 2021; Wherry *et al*., 2003). Variability in model outcomes is generated by randomization of this T cell exhaustion rate, which is selected from a shifted exponential distribution and sorted in increasing order as a function of pMHC reactivity (with a mean determined by model fitting, see Methods) (**Figure 1C**).

**Figure 1.**
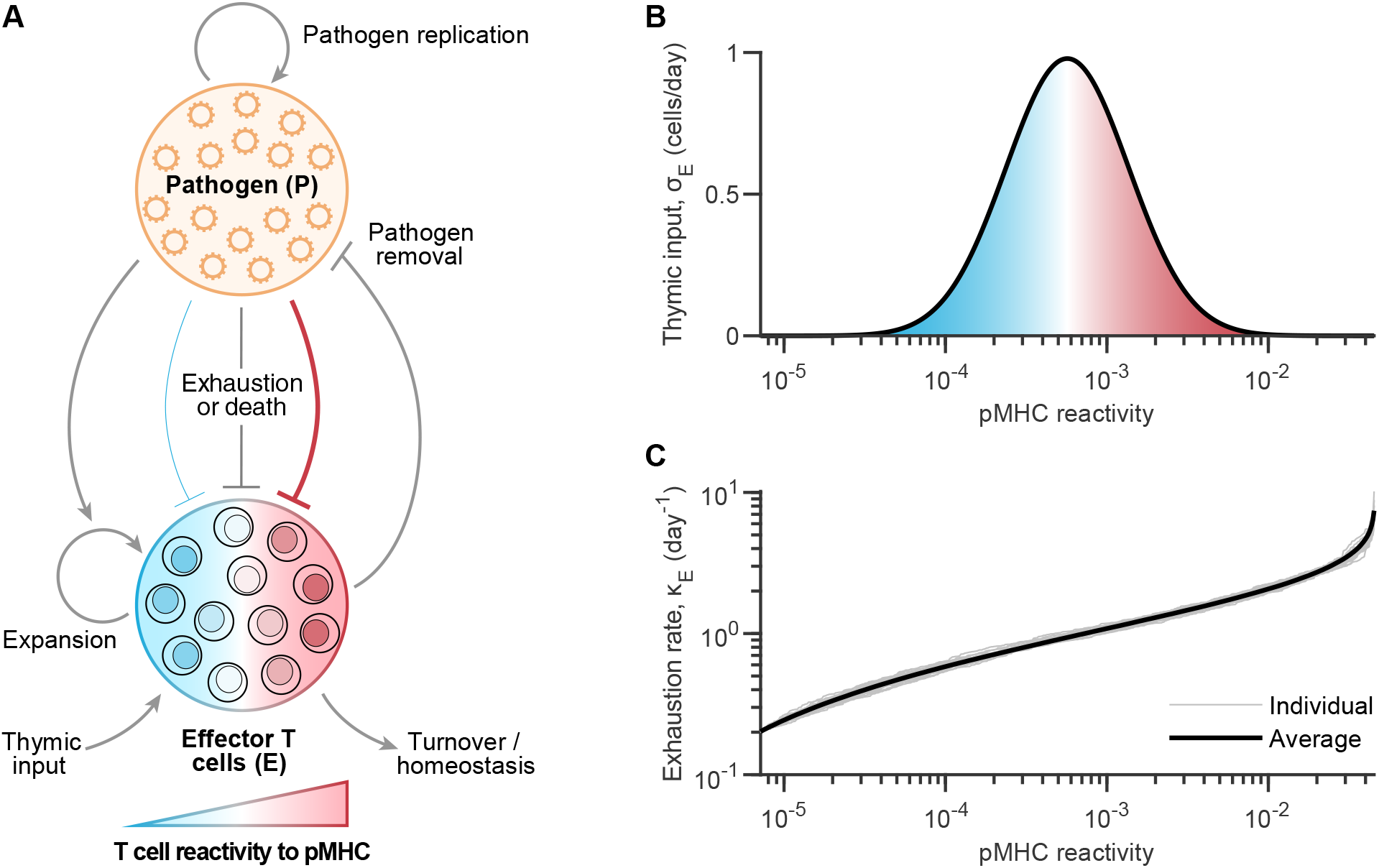
Implementing a computational population model defining the dynamics of pathogen replication and responding T cell clonotypes. A) Model scheme illustrating model variables and their interactions. The model is described by a system of integro-differential equations that govern the rates of change of the two key players, namely pathogen loads (*P*) and the effector T cells (*E*) that are specific for pathogen-derived antigens in the form of pMHC. The effector T cell population, encompassing the complete collection of pathogen-specific CD4^+^ and CD8^+^ T cells, are defined on a continuum according to their overall reactivity to pMHC (*a*_*k*_), i.e. strength of the overall TCR-pMHC interactions (the shade from blue to red represents the reactivity continuum of T cells to pMHC that ranges from low to high TCR affinities, respectively). Pathogen load is subject to replication as well as negative regulation by effector T cells. Effector T cells expand upon pathogenic exposure, with a constant low-level thymic input, natural turnover, and homeostatic competition. Pathogen persistence promotes T cell exhaustion and/or activation-induced cell death, with higher pMHC-reactivity T cells more susceptible to these than their low pMHC-reactivity counterparts. For simplicity, we (i) left out the role of other players such as antigen-presenting cells and B cells from the model in order to focus on the specific role of pMHC reactivity in defining dynamics, and (ii) did not explicitly distinguish between the action of CD4^+^ and CD8^+^ T cells. B,C) Functions depicting pMHC reactivity-dependent parameter values, wherein thymic input (B), σ_*E*_ = *f*(*a*_*k*_), mimics the shape of a theoretical log-normal distribution, and T cell exhaustion rate (C), *κ*_*E*_, is determined by sampling from a shifted exponential distribution of a set mean, and sorted in ascending order by assigning smaller depletion rates to T cells of low pMHC-reactivity, and *vice versa*.

Model parameters (**Table S1**) were obtained by fitting simulated pathogen loads to serum data of LCMV infections from Wherry *et al*. (Wherry *et al*., 2003) using a genetic algorithm (**Figure S1A** and **B**). Given that previous work showed LCMV-Cl13 replicates more rapidly in infected cells than LCMV-Arm (Bergthaler *et al*., 2010), we set the replication rate (denoted *r*_*P*_) for our simulated chronic viral infection to be higher than for the acute infection, while keeping all other parameters consistent between the two conditions. By modulating only the pathogen replication rate, we produced time courses that qualitatively matched both LCMV-Arm and LCMV-Cl13 replication dynamics (**Figures 2A, B**, and **S1A, B**). In our simulations, the acute viral load peaked at 5.8 days post infection, followed by rapid clearance around 6.4 days post infection based on 100 trial simulations (**Figures 2A** and **S1C**). In contrast, increasing the viral replication rate to simulate chronic viral replication resulted in prolonged infections with a median time to clearance of 61 days (based on 100 trial simulations) (**Figures 2B** and **S1C**). Thus, our model was able to reproduce pathogen loads characteristic of both acute and chronic infections by modifying only the rate of pathogen replication while keeping all other model parameters constant.

**Figure 2.**
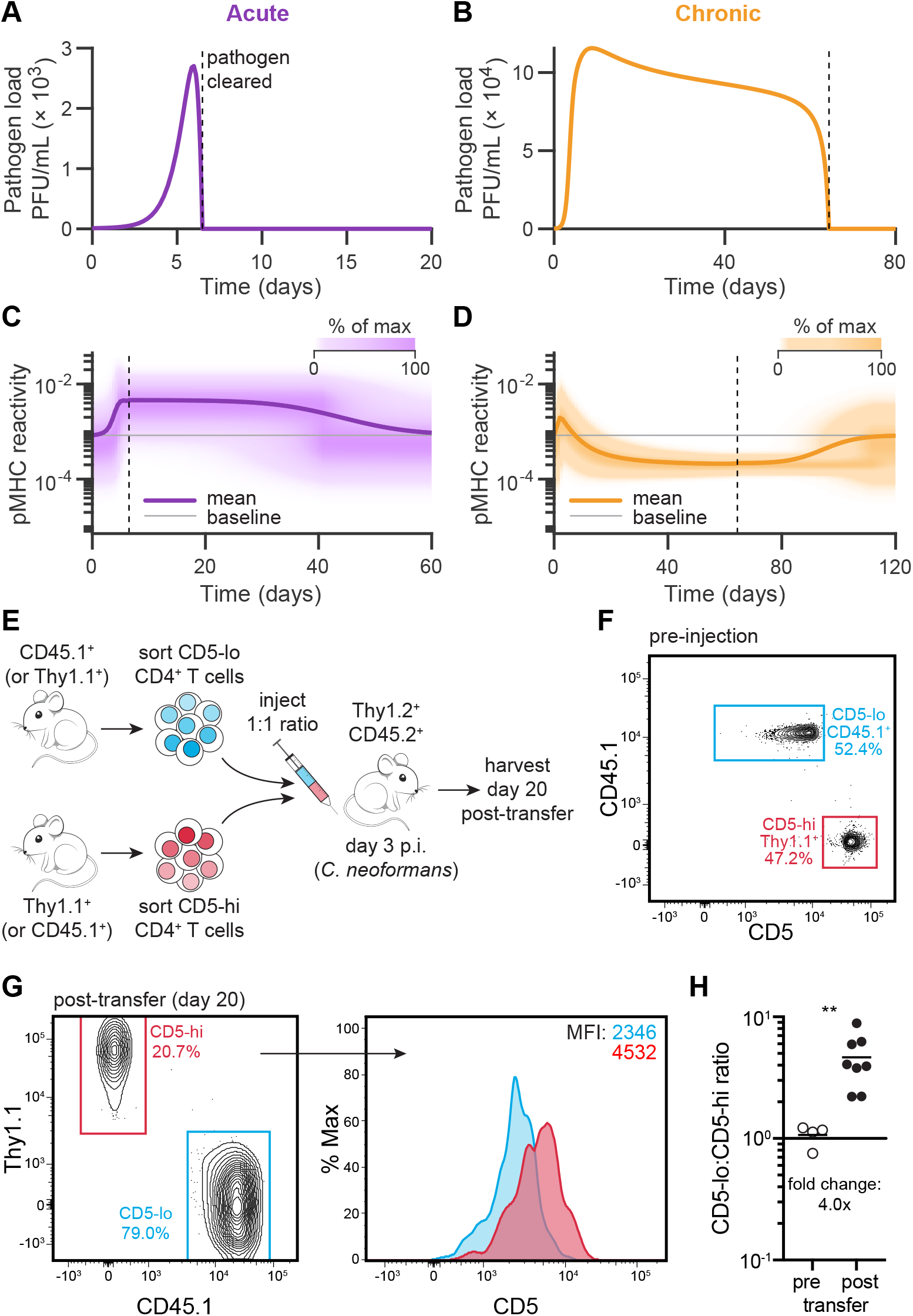
T cell pMHC reactivity evolution over time during a T cell response differs between acute and chronic infections. A-D) Simulations of pathogen loads and T cell levels in time during acute (A,C) or chronic (B,D) infection, showing time series simulations of pathogen loads (A,B) and heat maps representing the relative proportion of T cells across pMHC reactivities (C,D). Overlayed traces represent the mean pMHC-reactivity value in time, with the baseline value denoted in gray. Dotted black lines denote the time at which the pathogen is cleared. E) Schematic of experimental approach illustrating adoptive cell transfer of a 1:1 ratio of CD5^lo^ and CD5^hi^ naïve CD4^+^ T cells (either Thy1.1^+^ or CD45.1^+^) into congenic CD45.2^+^ Thy1.2^+^ recipient mice infected 3 days prior with chronic *C. neoformans*. F-G) Representative flow cytometry plot of CD5^lo^ CD45.1^+^ and CD5^hi^ Thy1.1^+^ transferred T cell populations, pre-injection into recipient infected mice (F), or 20 days post-transfer with CD5 expression levels shown as a histogram and mean fluorescent intensities (MFI) indicated in blue and red text (G). H) Ratio of transferred CD5^lo^ to CD5^hi^ T cells, pre-injection or 20 days post-transfer. **, *P* = 0.004 computed using a two-tailed Wilcoxon rank sum test, data from 2 independent experiments (n=8 mice).

### Chronic infection skews responding T cell clonotypes toward lower pMHC reactivities

Having developed our *in silico* model of acute and chronic pathogen infection, we next asked how the pMHC-reactivity profile of the antigen-specific T cell population evolved over time in each case. When we simulated effector T cell responses to acute infection, we found that the pMHC-reactivity profile of T cells shifted toward a higher pMHC-reactivity mean until peak pathogen replication was reached, followed by a gradual return to baseline after the infection was resolved (**Figure 2C** and **Movie 1**). Of note, the return of the pMHC-reactivity mean value to the pre-infection baseline was a result of omitting a memory T cell compartment from the model, since our focus was on the effector phase of the T cell response. The shift we observed in the pMHC-reactivity profile during acute infection agrees with previous experimental studies showing that T cells with greater pMHC reactivity expand to large numbers more readily upon antigenic stimulation (Busch and Pamer, 1999; King et al., 2012; Mandl et al., 2013; Rosenthal et al., 2012).

Next, we investigated whether the dynamics of pMHC reactivity among responding T cell clonotypes differed during chronic infection. In contrast to acute infection, the pMHC-reactivity distribution peaked during the early phases of chronic infection but was followed by a substantial shift toward lower pMHC reactivities, before returning to baseline after infection clearance (**Figure 2D**). This indicated that, as more T cells of high pMHC reactivity became functionally exhausted due to chronic antigen stimulation from persistent virus (and were thus removed from the active effector T cell pool), T cells of progressively lower reactivities to pMHC gradually predominated among responding T cell clonotypes (**Figure 2D** and **Movie 1**). Notably, this shift in pMHC reactivity seen over the course of a chronic infection, which results from a spectrum of TCR affinities present within the responding T cell population, is consistent with a previous report showing a skewing of the T cell response to persistent murine cytomegalovirus (MCMV) infection toward T cells with faster pMHC dissociation rates and thus lower binding affinities (Schober et al., 2020).

To further investigate whether our model was consistent with experimental data, we used a previously identified surface marker proxy for pMHC reactivity, CD5, whose expression levels on CD4^+^ T cells correlate with tetramer binding strength (Azzam et al., 1998; Mandl *et al*., 2013; Rogers et al., 2021). We sorted naïve CD4^+^ T cells on the 20% CD5^lo^ and CD5^hi^ cells, as previously described (Mandl *et al*., 2013; Rogers *et al*., 2021), mixed them 1:1 (identified by congenic markers, CD45.1 or Thy1.1), and adoptively transferred them to CD45.2^+^ Thy1.2^+^ recipient mice that were infected 3 days earlier with *Cryptococcus neoformans*, a persistent pulmonary fungal pathogen (Schneider et al., 2020) (**Figures 2E, F** and **S1D, E**). In contrast to what was previously described during acute infections, where the CD5^hi^ CD4^+^ T cells predominated the response on day 8 post infection (Mandl *et al*., 2013), in the later stage of the anti-*Cryptococcus* CD4^+^ T cell response in the lung, the CD5^lo^ CD4^+^ T cells outnumbered CD5^hi^ CD4^+^ T cells 4-fold (**Figure 2G, H**). Taken together, these results suggest that the temporal evolution of pMHC reactivities of T cells contributing to the response to an acute compared to a chronic infection is distinct, with T cells of lower pMHC reactivities predominating during the chronic infection phase, whereas T cells of greater pMHC reactivity predominate during an acute infection.

### The distribution of pMHC reactivities of responding T cells impacts time to infection clearance

Having established that the pMHC reactivity of effector T cells changed differentially in acute versus chronic infection, we wanted to determine whether altering the pMHC-reactivity profile of the antigen-specific T cell population would lead to changes in the duration of pathogen replication upon infection using our simulations. To do so, we targeted model parameters affecting either the mode or the span of the T cell pMHC-reactivity distribution. We used the time at which pathogen burden returned to zero post infection as a measure to assess: (i) class of infection (i.e., acute vs. chronic), and (ii) effectiveness of the T cell response in clearing infection. Gradually varying the mode of the pMHC-reactivity profile over 40 log-linearly spaced values between 10^−4^ and 10^−2^ and running 50 trial simulations per mode value revealed three distinct clusters with different durations of infection (**Figure 3A**). When the mode of antigen-specific pMHC-reactivities of the T cell repertoire was too low (< 2 × 10^−4^), the infection persisted indefinitely, indicating that T cells failed to clear the pathogen. At intermediate values of pMHC reactivity (between 2 × 10^−4^ and 10^−3^), the time to clearance was around 100 days, comparable to chronic LCMV-Cl13 infection. When reactivity to foreign pMHC was high, the infection was cleared within 10 days, representative of an acute infection (**Figure 3A**). Varying the standard deviation of the pMHC reactivity distribution also affected pathogen replication kinetics, with a larger span of the pMHC reactivity promoting more rapid pathogen clearance over T cell repertoires with narrower affinity distributions (**Figure 3B**).

**Figure 3.**
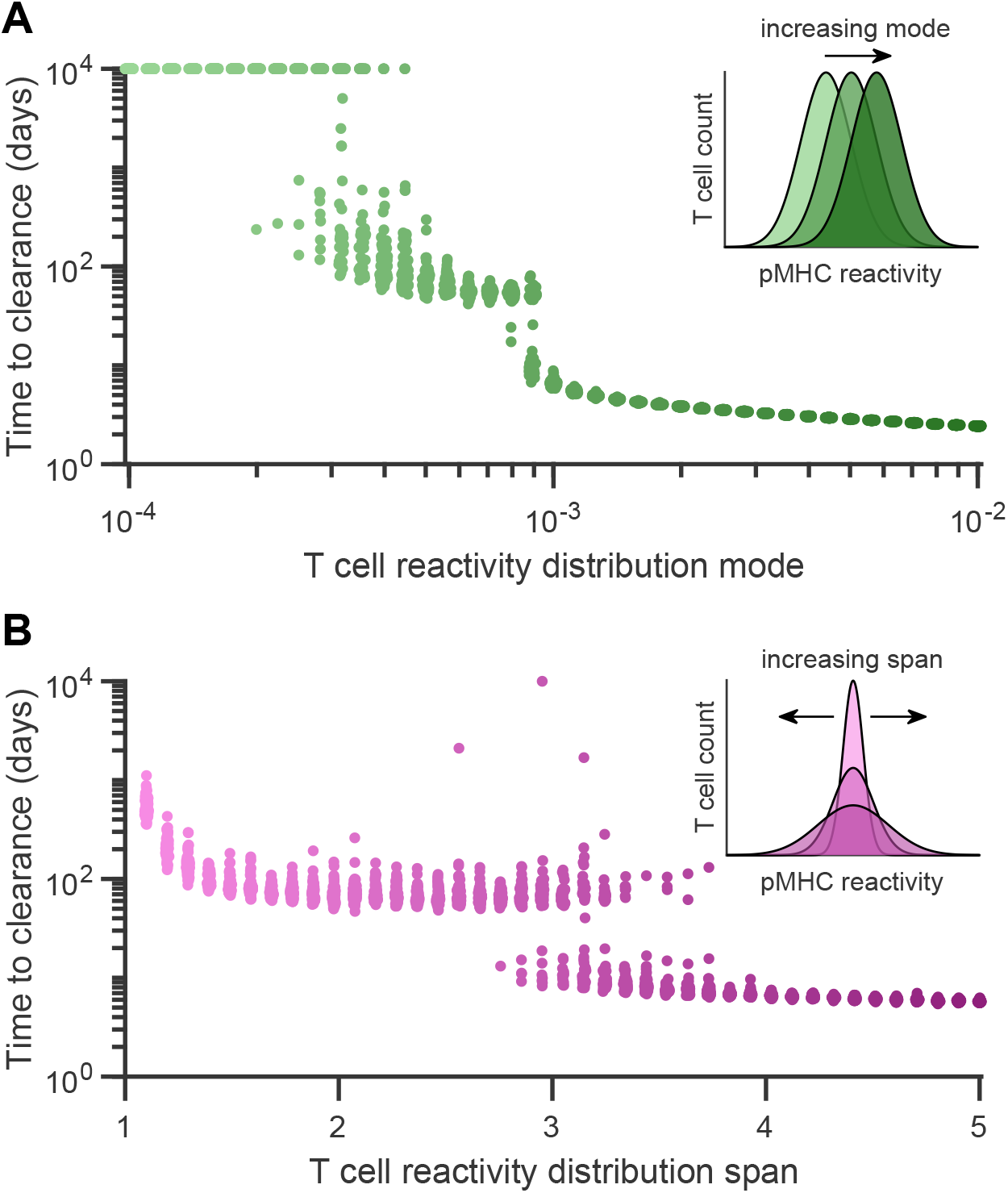
Mode and span of the naïve T cell pMHC-reactivity profile impacts pathogen clearance times. A-B) Time required to clear a replicating pathogen *in silico* when varying A) the mode, and B) the span of the initial T cell count as a function of pMHC reactivity. Each data point shows the result of a single simulation trial for a given, randomized set of parameters for T cell exhaustion (*κ*_*E*_) as determined by the relation defined in Figure 1C. Insets indicate how the T cell count of the starting (pathogen-free/baseline) configuration, as a function of pMHC reactivity, is altered by increasing its mode (A) or span (B).

Taken together, our results demonstrate that the pMHC-reactivity profile of responding T cells during infection has a pronounced effect on determining infection duration, with outcomes ranging from rapid pathogen clearance to the complete failure of the T cell response to resolve the infection. Of note, rather than observing gradual shifts in the time to clearance with *in silico* manipulations of the pMHC-reactivity profile, we found sharp jumps between different infection clearance times (namely from acute to chronic, and chronic to indefinitely persistent). By reducing the model to a one-clone, 2-dimensional system of ordinary differential equations and performing a bifurcation analysis (**Figure S2**, and Supplementary Text), we found that distinct solution trajectories through state space are responsible for producing separate clusters of infection durations. Importantly, analysis of the one-clone model showed that, exclusively in the case of chronic pathogen load, the evolution of the T cell profile toward lower pMHC reactivities is responsible for the eventual resolution of chronic infection (**Figure S2** and **Movie 2**).

### Absence of T cells with low pMHC-reactivity generated by N-nucleotide diversity delays the clearance of chronic infection

Our results thus far showed that an increased contribution from T cell clonotypes with low pMHC reactivity is vital in clearing chronic, but not acute, infections. Thus, we next investigated whether populating a TCR repertoire exclusively with higher pMHC reactivity T cells would affect the clearance of either an acute or a chronic infection *in silico*. We tackled this question in two ways. First, in our simulations, we removed T cells with lower pMHC reactivity by eliminating all T cells possessing a pMHC reactivity value below a predefined cut-off threshold without altering the total number of responding T cells, and then progressively increased this threshold to remove up to half of the pMHC reactivity distribution (**Figure 4A**). Our simulations demonstrated that impeding the shift toward lower TCR affinities during a chronic infection by progressively removing T cells with lower pMHC reactivity in this manner dramatically prolonged the time to clearance for the chronic infection cluster (**Figures 4B**, and **S3**).

**Figure 4.**
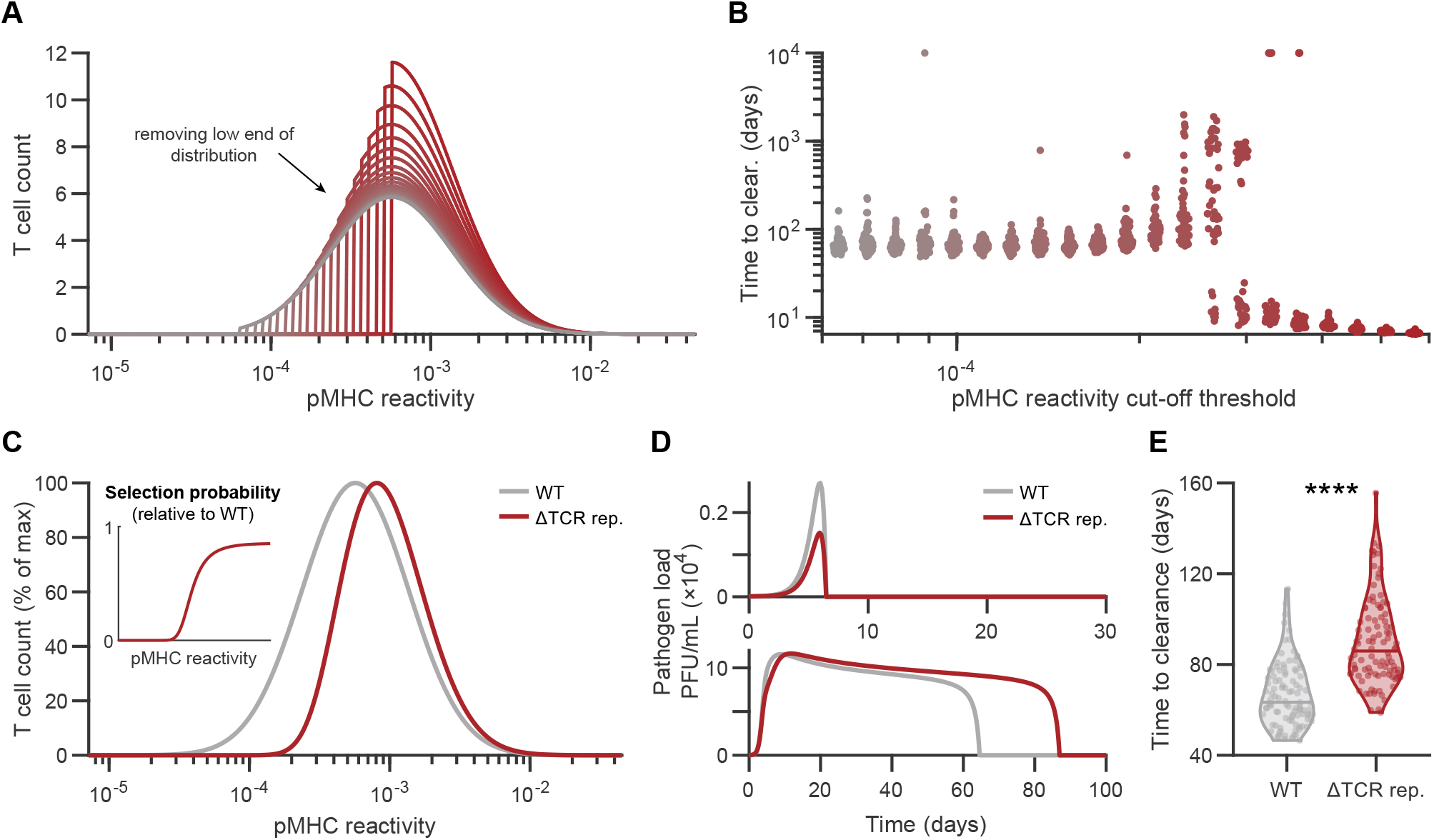
Deficiency in TCRs with low pMHC-reactivity *in silico* leads to prolonged chronic, but not acute, pathogen replication. A) Distributions of T cells as a function of pMHC reactivity obtained by successively removing low-affinity T cells using different cut-off thresholds from the model’s starting configuration, while keeping the total number of T cells conserved, until only the upper half of the distribution remained. These starting configurations were generated by setting all values of the thymus input, σ_*E*_, for T cells below a given pMHC-reactivity threshold to 0. B) Time to pathogen clearance as a result of increasing the cut-off threshold for T cell reactivity to pMHC corresponding to the configurations shown in A. For each cut-off threshold, 50 simulation trials were performed as described in Fig. 3. C) Theoretical T cell pMHC-reactivity configuration of an altered TCR repertoire (denoted ΔTCR repertoire) deficient in low-affinity T cells relative to the WT configuration. The ΔTCR repertoire was assumed to have a lower thymic selection probability (relative to WT repertoire), implying a lower value of the thymic input parameter, σ_*E*_, for T cells with low pMHC reactivity. Inset: Probability of selection, relative to the WT repertoire, for T cells of the ΔTCR repertoire, with fewer T cells of low reactivity to pMHC being sourced by thymus selection. D) Model simulations comparing representative pathogen load traces of WT (gray) or ΔTCR repertoire (red) systems during acute (top) or chronic (bottom) infection. E) Time to clearance of chronic infections for 100 model simulations from WT and ΔTCR repertoire systems. ****, *P* = 1.51×10^−17^ computed using the Wilcoxon rank sum test.

Interestingly, this occurred only up to a certain cut-off threshold (no T cells with pMHC reactivity below *a*_*k*_ ≈ 3 × 10^−4^), beyond which the greater number of only high pMHC reactivity T cells prevented infection chronicity altogether because of effective early pathogen clearance, leading to the formation of an acute infection cluster (**Figure 4B**). When the number of high pMHC reactivity T cells was kept unchanged, the acute cluster did not form (**Figure S3**).

While gradually removing all effector T cells below a certain pMHC-reactivity threshold allowed us to investigate the distinct roles of T cells with low versus high pMHC reactivity, we next designed a second approach in which we also modified our model to simulate a more biologically plausible change in the pMHC reactivity profile. In this approach, we defined an altered TCR repertoire (ΔTCR rep.), wherein the probability that low-reactivity T cells selected for in the thymus was reduced, while the selection probability of high-reactivity T cells was left relatively unchanged (**Figure 4C**). We found that this ΔTCR rep. did not alter replication kinetics of an acute pathogen (**Figure 4D**, top). However, testing the ΔTCR rep. model with a chronic pathogen revealed that chronic infection clearance was impaired (**Figure 4D**, bottom), and time to clearance was significantly longer (**Figure 4E**). In summary, altering the pMHC-reactivity profiles of responding T cells by introducing reductions in T cells with low pMHC-reactivity showed that modulating only the TCR repertoire pMHC reactivity led to impaired control of chronic, but not acute, infections.

The proposed hypothesis that N-diversity mediated by TdT, which accounts for 90-95% of the TCR repertoire diversity (Cabaniols *et al*., 2001), disproportionately generates lower-affinity TCRs (Vrisekoop *et al*., 2014) has been difficult to address experimentally without a specific prediction of the type of infection that these low-affinity TCRs are important for with regard to curtailing pathogen replication. Based on our modeling results suggesting that a TCR repertoire deficient in T cells with low pMHC reactivity would lead to an impaired effector T cell response during chronic infection, we next sought to test our model prediction *in vivo* in mice and ask whether TdT might benefit the host by promoting the shift of T cells from high- to lower-pMHC reactivity. Given that TdT also inserts non-templated nucleotides into the B cell receptor during B cell development (Jackson et al., 2013), and the altered B cell receptor repertoire might therefore impact viral clearance, we restricted the TdT-deficiency to the T cell compartment only, with B cells expressing normal levels of TdT. To do so, we generated bone marrow chimeras (**Figure 5A**) whereby TCRβ KO bone marrow was mixed 1:1 with either WT bone marrow (leading to development of WT B and T cells), or with bone marrow obtained from TdT and JH double KO mice (leading to the development of WT B cells and TdT KO T cells). We verified that our bone marrow reconstitutions led to a 1:1 ratio of hematopoietic cell development from each of the donors, identified using congenic markers (**Figure S5A**). In line with previous results from full TdT KO mice in response to acute LCMV infection (Gilfillan *et al*., 1995a), we observed no differences in the viral load following LCMV-Arm infection between the WT T cell and TdT KO T cell groups, both 2 days post infection in the serum, and 6 days post infection in the total spleen homogenate (**Figure 5B**). In contrast, mice with TdT KO T cells had a significantly higher viral load during the chronic phase (day 40-45) following infection with LCMV-Cl13 (**Figure 5C**). Since irradiated bone marrow recipients still retain some endogenous hematopoietic stem cells, we repeated the above experiment using TCRβ KO recipient mice instead (i.e., mice lacking endogenous αβT cells), and obtained similar results (**Figure S4B** and **S4C**). Taken together, our modeling predictions in conjunction with our experimental results support the hypothesis that TdT benefits hosts challenged with chronic infections, leading to faster viral clearance rates.

**Figure 5.**
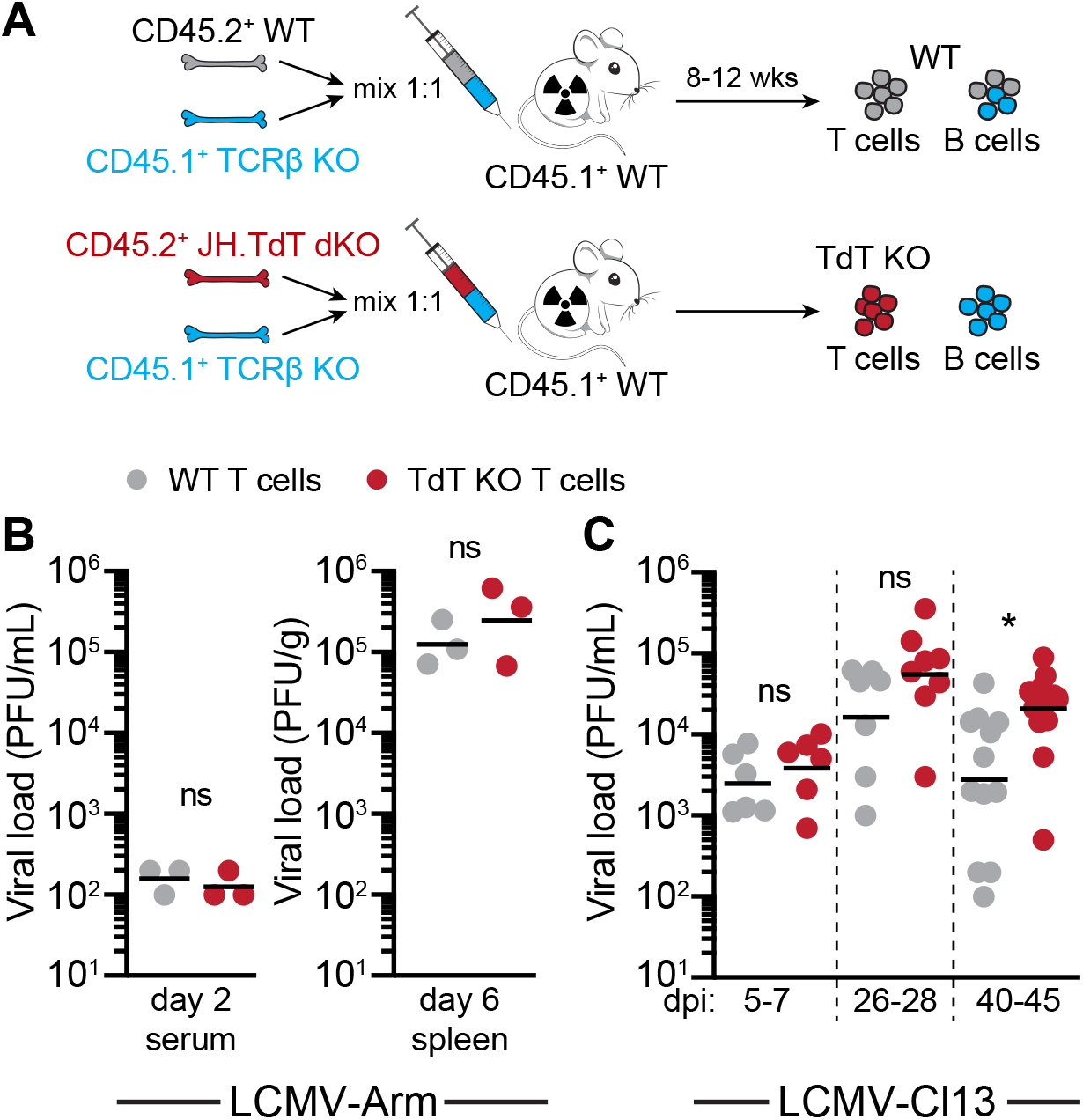
Loss of TdT restricted to T cells *in vivo* impairs control of chronic, but not acute, infections. A) Generation of bone marrow chimeric mice possessing WT B cells, and either WT T cells (reconstitution of irradiated mice with 1:1 ratio of bone marrow from WT mice and TCRβ^−/–^ mice) or TdT KO T cells (reconstitution with 1:1 ratio of bone marrow from JH and TdT double KO mice and TCRβ^−/–^ mice). B) LCMV-Arm viral loads in the serum (left) and spleen (right) of mice with WT or TdT KO T cells measured at 2 or 6 days post-infection, respectively (n=3 mice per group). C) LCMV-Cl13 viral loads in the serum of mice with WT or TdT KO T cells in early (5-7 days post-infection), mid (26-28 days post-infection) or late (40-45 days post-infection) stages of viral replication (n = 6-18 mice per group). *, *P* = 0.013 computed using a Kruskal-Wallis test.

## Discussion

N-nucleotides added by TdT during V(D)J gene segment recombination contribute enormously to the diversification of the TCR repertoire (Cabaniols *et al*., 2001). Yet, despite the fact that TdT is found in all jawed vertebrates with adaptive immune systems studied thus far (Litman *et al*., 2010), the specific contexts in which these non-germline TCRs are better poised to control pathogen replication have not been clear (Gilfillan *et al*., 1995a; Gilfillan *et al*., 1995b). Here we combined computational modelling and experimental approaches to investigate the temporal evolution of pMHC reactivities of responding T cells during infection, and its impact on pathogen clearance. We developed a computational model that (1) produced time courses characteristic of infections with both acute and chronic pathogens, and (2) incorporated a continuum-affinity formalism to track T cell pMHC-reactivity distributions over time. Using the *in silico* model we developed, we made two predictions that we tested experimentally. First, we showed that, while in acute infection T cells with high pMHC reactivity predominate (Bachmann et al., 1997; Busch and Pamer, 1999; McHeyzer-Williams and Davis, 1995), during chronic infection T cells with low pMHC reactivity contribute disproportionately. Second, we found that the removal of low pMHC-reactivity T cells leads to a delay in chronic, but not acute, pathogen clearance in the model, which we replicated in infected mice when T cells were TdT deficient. Importantly, our data corroborate prior experimental work showing no differences in clearance by TdT KO mice of the acute viral pathogens Vesicular Stomatitis, Sendai, Influenza A, and LCMV-WE (Gilfillan *et al*., 1995a; Haeryfar *et al*., 2008). Thus, while it has been proposed that a TdT-deficient TCR repertoire may have specific ‘holes’ with regard to antigen specificities represented, this has so far not been supported by experimental evidence. Indeed, without accounting for possible differences in T cell clonotype precursor frequencies, our model predicts longer times to clearance for chronic pathogens. Our work therefore suggests a hitherto undescribed benefit for TCR repertoire diversification by TdT in chronic infection control.

A TdT-deficient repertoire and its consequences for pathogen control are relevant not only for understanding the broad evolutionary conservation of TdT across vertebrates, but also in the context of neonatal immunity, given that the TCR repertoire is initially generated in the absence of TdT. TdT expression is first detected in thymocytes in mice and humans 3-5 days and 20 weeks after birth, respectively (Bogue et al., 1992; Bonati et al., 1994). Lacking N-nucleotide additions, neonatal TCR sequences are shorter, are more likely to be shared between individuals (public) (Yassai et al., 2002; Yassai and Gorski, 2000) and are more cross-reactive (Gavin and Bevan, 1995). Interestingly, in line with TdT KO T cells having greater pMHC reactivity, it has been shown that the neonatal repertoire is more self-reactive due to a greater affinity for pMHC, and neonatal T cells more prone to tolerance (Rudd, 2020). To what extent the neonatal TCR repertoire versus other epigenetic or transcriptional differences described in neonatal compared to adult T cells play a role in altered responses to infection requires further analysis.

Although our model was fit to viral replication data from acute and chronic strains of LCMV (Wherry *et al*., 2003), its conclusions may be generalizable to other pathogens. For instance, we found that following infection with the pulmonary fungal pathogen, *C. neoformans*, CD4^+^ T cells with low self-pMHC reactivity, and thus low foreign reactivity (Mandl *et al*., 2013), predominated among the responding effector T cells during the chronic infection phase. These experimental findings expand on previous work that suggested that, during chronic MCMV infection, the T cell repertoire shifts towards cells with lower pMHC binding strength (Schober *et al*., 2020). Notably, in our current model we did not distinguish between CD4^+^ and CD8^+^ T cell responses, and it is possible that there are key differences between these T cell subsets with regard to the role of TdT and functional biases among TdT-generated TCRs, which need to be further explored.

Moreover, our model did not consider memory T cells following pathogen clearance, and experimental data suggests that memory T cells are differentially selected for in terms of their reactivity to pMHC, resulting in a final steady-state pMHC-reactivity distribution of memory T cells that differs from the starting naïve T cell pMHC reactivity distribution (Andargachew *et al*., 2018; Busch and Pamer, 1999; Mandl *et al*., 2013).

Our work revealed the effects of varying features of the pMHC-reactivity distribution of responding T cells on pathogen clearance and suggested a differential role between T cells with low- and high-reactivity to pMHC during different phases of the immune response. This is particularly intriguing, given recent observations that in both mice and humans, T cells with lower self-pMHC reactivity (low CD5 surface levels) express higher levels of *Dntt*, the gene encoding TdT (Fulton et al., 2015; Rogers *et al*., 2021; Sood et al., 2021). Thus, differences in TdT expression level during development in individual thymocytes may ultimately contribute to the numbers of N-nucleotides inserted into the recombining TCR and be a critical variable impacting strength of pMHC reactivity. To provide additional insight into whether there are indeed different roles for TdT-mediated versus germline-encoded TCRs during infection, comprehensive TCR sequencing studies will be a powerful tool. Overall, our model formalism provides a foundation for further studies of T cell pMHC-reactivity distributions over the course of an immune response, and it will be particularly interesting to investigate whether, as our model suggests, TdT-dependent TCRs are important in the control of other chronic pathogens and are perhaps making underappreciated contributions in settings such as cancer and autoimmunity.

## Acknowledgments

This work was done in Tiohtiá:ke/Montreal on the traditional territory of the Kanien’kehà:ka, a place which has long served as a site of meeting and exchange amongst many First Nations including the Kanien’kehà:ka of the Haudenosaunee Confederacy, Huron/Wendat, Abenaki, and Anishinaabeg. We honour, recognize, and respect these nations as the traditional stewards of these lands and waters. We would like to thank the animal facility staff at McGill University for their excellent care of our animal colony, and C. Stegen and J. Leconte at the Cell Vision Core Facility for cell sorting, and P. Artusa for initial experiments at an earlier stage of the project. H.J. was supported by a Post-Graduate Doctoral Award (NSERC) and a B2X Doctoral Award (FRQNT). D.R. was supported by a Frederick Banting and Charles Best Canada Graduate Doctoral Award (CIHR) and a Tomlinson Doctoral Fellowship (McGill). J.N.M. is a Canada Research Chair for Immune Cell Dynamics. This work was supported by NSERC (Discovery Grants #2019-04520 to A.K. and #2016-03808 to J.N.M.).

## Author Contributions

J.N.M. and A.K. conceived the study, supervised, and led the study implementation, and acquired the financial support for this project. H.J. designed and analysed the computational model and wrote the manuscript with input from D.R., J.N.M., and A.K.

D.R. designed, performed, and analysed experiments, with key experimental input from G.P. H.J. and D.R. prepared the figures.

## Figure Legends

**Movie 1** (Related to Figure 2). Time series simulations of the model during acute (A-C) or chronic (D-F) infection, showing pathogen loads (A,D), heat maps representing the relative proportion of T cells across pMHC reactivities (B,E), and evolution of T cell proportions as a function of pMHC reactivity at each time point (C,F). Overlayed traces in B and E represent the mean reactivity value in time, weighted by the proportion of T cells of given reactivity values. Dotted black lines denote the time at which the pathogen is cleared.

**Movie 2** (Related to Figure 3). Nullclines of the reduced, 2-dimensional, one-clone model plotted in logarithmic (A) and linear (B) scales over time during chronic pathogen replication. Nullclines vary over time since the parameters for pMHC reactivity (*a*_*k*_) and exhaustion rate (*κ*_*E*_) of the one-clone model were set to their respective weighted average values across all T cells of the full model at each time point. Given the relative proportions of T cells of different pMHC reactivity values vary through time in the full model, the weighted average of these parameters also changes over time. Black lines denote nullclines of the pathogen load (*P*) nullclines while the gray line denotes the effector T cell (*E*) nullcline. Superimposed in green is the pathogen load and the total T cell count (across all values of pMHC reactivity) simulated from the full model across time. Refer to Supplementary Text for more details.

## Materials and Methods

### Mice

C57BL/6, congenic CD45.1^+^, congenic Thy1.1^+^, and TCRβ^-/-^ mice (Mombaerts et al., 1992) were purchased from Jackson Laboratories (Bar Harbor, ME). The TdT^-/-^ mice were shared by Dr. A. Feeney (The Scripps Research Institute) (Gilfillan et al., 1993) and the JH^-/-^ mice were shared by Dr. J. Fritz (McGill) (Gu et al., 1993). All mice were on a C57BL/6 background, bred in-house and experiments performed at 6-12 weeks of age with both males and females. Animal housing, care, and research were in accordance with the Guide for the Care and Use of Laboratory Animals and all procedures performed were approved by the McGill University Animal Care Committee.

### Pathogens stocks and infections

*LCMV*. LCMV-Arm and -Cl13 strains were propagated from stocks provided by Dr. M. Richer (University of Indiana) on BHK-21 or L929 cells (ATCC). Briefly, virus was added at MOI 0.01, incubated for 90 minutes in serum-free media at 37°C in 5% CO_2_, then topped up with complete media for incubation for another 48 hours before harvesting the supernatant. BHK-21 cells were cultured in EMEM supplemented with 0.1% penicillin/streptomycin, 1% L-glutamine, 1% non-essential amino acids, 1% sodium pyruvate, and 10% FBS and maintained at 37°C in 5% CO_2_. L929 cells were cultured in RPMI supplemented with 10% FBS, 1% L-glutamine, and 1% penicillin/streptomycin.

Mice were infected with 2×10^5^ plaque forming units (PFU) of LCMV-Arm by intra-peritoneal injection or 2×10^6^ PFU by intravenous injection for LCMV-Cl13 as previously described (Richer et al., 2013; Wherry *et al*., 2003). Mice were bled by either tail artery or cardiac puncture into sterile eppendorf tubes kept on ice, blood was spun down at 12,000 rpm for 10 minutes and serum aliquoted and frozen for viral titer determination.

Spleens were collected in 1% RPMI and weighed. Spleens were placed in Lysing Matrix D tubes (MP Biomedicals) and homogenized with a MagNA Lyser (Roche) at 6000 rpm for 40 seconds. Spleen homogenate was then spun down at 12,000 rpm for 10 minutes at 4°C and supernatant was transferred to a separate sterile tube and re-spun at 12,000 rpm for 10 minutes at 4°C then aliquoted and frozen for viral titer determination. Viral titers (stocks used, mouse serum, and tissue samples) were determined by plaque assay with Vero cells (Ahmed *et al*., 1984). Briefly, Vero cell monolayers were infected with 100 μL of serially diluted serum (1 in 10 dilutions from 10^−1^ to 10^−7^) and incubated for 90 minutes at 37°C in 5% CO_2_. Infected cells were then overlaid with 1% agarose (Wisent) and incubated for 3 days at 37°C in 5% CO_2_. A second agarose overlay supplemented with 1% neutral red was then added and cells incubated for 24 hours at 37°C in 5% CO_2_, after which plaques were counted.

*C. neoformans*. The H99 strain was provided by K. Kwon-Chung (NIH). Frozen stocks (−80°C) were prepared in 15% glycerol from fresh cultures from a YPD agar plate. Three days before infection, *C. neoformans* was scraped from the frozen stock and streaked onto a YPD agar plate. One day prior to infection, a single colony was inoculated and incubated for 12-16 hours at 30°C with continuous agitation in YPD broth. Immediately before infection, C. neoformans was resuspended in cold PBS. Mice were then anesthetized with isoflurane and infected by intrapharyngeal aspiration with 5×10^3^ colony forming units (CFU) in 20 μL of PBS. Mice were sacrificed and tissue collected 23 days after infection. *C. neoformans* CFUs were determined as previously described (Schneider *et al*., 2020).

### Lymphocyte isolation

For the *C. neoformans* infections, prior to harvest an intravascular stain using 2.5 μg anti-CD45 (30F11) was performed as previously described (Anderson et al., 2014). Infected lungs were harvested in cold PBS and minced with scissors. Lung was then digested at 37°C with agitation for 30 minutes in digestion buffer (1 mg/mL collagenase D, 50 U/mL DNase I, 1 mg/mL hyaluronidase, 1% L-glutamine, 1% pen/strep in RPMI). Tissue was then passed through a 100-μm filter with PBS supplemented with 1% FBS and resuspended in 10mL of 37% Percoll in RPMI. Samples were centrifuged at 3000 rpm for 20 minutes at 22°C. ACK lysis buffer (Life Technologies) was added for 3 minutes, samples washed with PBS, refiltered, and resuspended in complete RPMI (10% FBS, 1% L-glutamine, 1% HEPES buffer, 1% pen/strep, 1% sodium pyruvate, 1% non-essential amino acids, 0.1% 2-mercapto-ethanol 1000X solution). Dilution of single-cell suspensions at 1:10 in Trypan Blue and manual counting of live cells (Trypan Blue-negative) on a hemacytometer was used to determine total cell counts.

Spleen and peripheral lymph nodes (inguinal, axillary, brachial, and mesenteric) were collected and passed through a 70-μm filter with 1% RPMI (1% penicillin/streptomycin, 1% L-glutamine, and 1% FBS). ACK lysis buffer (Life Technologies) was added for 3 minutes, samples washed with PBS, refiltered and resuspended in 1% RPMI. Dilution of single-cell suspensions at 1:10 in Trypan Blue and manual counting of live cells (Trypan Blue-negative) on a hemacytometer was used to determine total cell counts.

### Bone marrow chimeras

Bone marrow was collected from the femurs and tibias of donor mice (either JH^-/-^, TdT^-/-^, TCRβ^-/-^, or B6 WT) by flushing the marrow from the bones with cold 1% RPMI. Bone marrow cells were then passed through a 70-μm filter with 1% RPMI and red blood cells lysed with ACK lysis buffer (Life Technologies) and cell counts determined as above. Recipient mice (either B6 CD45.1^+^ or TCRβ^-/-^ CD45.1^+^) were irradiated twice at 550 rads 3 hours apart and reconstituted with a 1:1 mix of 2.5×10^6^ cells per genotype that were injected i.v. within 5 hours of the first irradiation. To establish the WT chimera (WT T cells and B cells) B6 and TCRβ^-/-^ bone marrow cells were mixed at equal proportions; to make the T cell restricted TdT^-/-^ chimeras (WT B cells) bone marrow cells from JH^-/-^ TdT^-/-^ mice were mixed 1:1 with TCRβ^-/-^ bone marrow cells. Recipient mice were given neomycin water (2g/L) 2 days prior to bone marrow transfer and kept on the antibiotic water for 2 weeks following transfer. Mice were used 8-12 weeks post irradiation and bone marrow reconstitution.

### Flow cytometry

Samples were incubated in Fixable Viability Dye (AF780, Life Technologies) diluted in PBS for 20 minutes at 4°C. Extracellular antibodies were diluted in FACS buffer (2% FBS and 5mM EDTA in PBS) with Fc Block (Life Technologies) and incubated for 30 minutes at 4°C. For intracellular staining, samples were then incubated in FoxP3 Transcription Factor Fixation/Permeabilization Concentrate and Diluent (Life Technologies) for 30 minutes at 4°C. Intracellular antibodies were diluted in Permeabilization Wash Buffer (Life Technologies) and samples were incubated for 30-60 minutes at 4°C. Directly conjugated antibodies used were as follows: TCRb (H57-597), CD4 (RM4.5), CD8a (53-6.7), CD5 (53-7.3), Foxp3 (FJK-16 s), CD44 (IM7), CD62L (MEL-14), CD25 (PC61.5), CD45.1 (A20), CD45.2 (104), PD-1 (29F.1A12), B220 (RA3-6B2), NK1.1 (PK126). For all flow cytometry experiments, cells were acquired using an LSRFortessa (BD Bioscience) and analyzed with FlowJo software (BD Bioscience).

### Cell sorts

Cell sorts were performed as previously described (Rogers *et al*., 2021). Briefly, lymphocytes from Thy1.1^+^ or CD45.1^+^ congenic mice were isolated in single cell suspension as described. Spleens and lymph nodes (inguinal, axially, brachial, mesenteric, and cervical) were pooled from 15 mice for each congenic marker. Cells were then magnetically enriched for total CD4^+^ T cells (Stemcell EasySep CD4^+^ T cell Enrichment kit or Miltenyi Biotec CD4^+^ T cell Isolation Kit). Enriched CD4^+^ T cells were stained with surface antibodies for 1 hour at 4°C. Naive CD4^+^ T cells were sorted on singlets, CD4^+^, CD8^-^, CD62L^hi^, CD44^lo^, and 20% CD5^lo^ or CD5^hi^. Sorts were performed on a FACS Aria III (BD Bioscience). All cell populations were sorted to >90% purity.

### Adoptive cell transfers

*C. neoformans* infection. All donors and recipients were sex matched. 15 CD45.1^+^ or Thy1.1^+^ mice were used as donors to obtain a total of 8-14 × 10^6^ cells for each of 20% CD5^lo^ and 20% CD5^hi^ cells sorted as described above. Sorted CD5^lo^ and CD5^hi^ were then mixed in a 1:1 ratio. 4-7×10^6^ of each sorted population was adoptively transferred into CD45.2^+^ Thy1.2^+^ recipients that were infected with 5×10^3^ CFU of *C. neoformans* 3 days prior to transfer. Cells were isolated from the lungs of recipient mice 20 days post-transfer.

### Statistical analyses of experimental data

Group comparisons were performed using Prism V9 (GraphPad). The cut-off for significance considered was p<0.05. Information about implemented statistical tests and sample sizes for individual experiments is provided in the figure legends.

### Computational modelling

To study a continuum of antigen-specific T cell affinities in the context of acute vs. chronic pathogen infections, we used the following system of integro-differential equations based on Fig. 1:

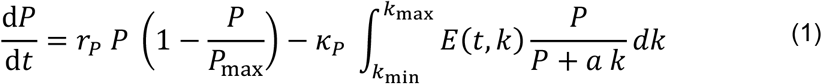

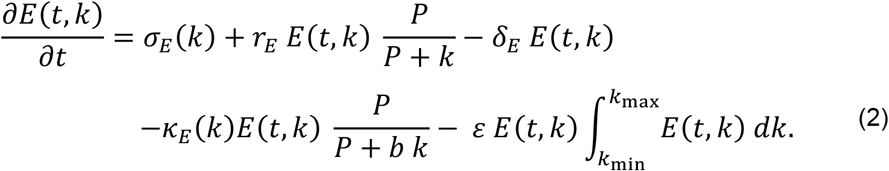

This was implemented in a manner similar to the model presented in (Jaberi-Douraki et al., 2015). In Eqs. 1 and 2, *P*(*t*) represents the pathogen load in time, and *E*(*t, k*) represents the time-dependent reactivity-continuum of effector T cells, with T cell pMHC reactivity taken to be proportional to the quantity 1/*k*, where *k* is the pathogen load for half-maximum activation of T cells. To facilitate understanding of pMHC-reactivity distributions, we defined a new, unitless quantity *a*_*k*_ (whose magnitude is equivalent to 1/*k*) as a “measure” for T cell reactivity to pMHC (Standifer *et al*., 2009). While this quantity is related to the overall binding avidity between T cells and APCs, we considered only affinity-related changes in TCR binding strength, and thus used the term pMHC reactivity to avoid confusion with other factors that can affect binding avidity. For simplicity, in Eq. 1 we assumed a logistic growth of the pathogen (with replication rate *r*_*P*_ and carrying capacity *P*_max_) as was done in previous models of LCMV infection (Bocharov, 1998; Keşmir and De Boer, 2003). Clearance of the pathogen by T cells was described by a product of the T cell number (*E*) and an increasing first-order Hill function of pathogen load with a maximum rate *κ*_*P*_, and a half-maximum pathogen load *a*. *k* (where *a* is a scaling factor). We impose the condition that the pathogen replication rate (*r*_*P*_) for chronic pathogens to be larger than that for acute pathogens, as suggested previously in the case of LCMV-Cl13 vs. LCMV-Arm, respectively (Bergthaler *et al*., 2010; Sullivan et al., 2011).

The terms included in effector cell dynamics of Eq. 2 were pMHC reactivity-dependent thymic input (σ_*E*_(*k*)), T cell replication that depends on pathogen load according to a first-order Hill function (with maximum replication rate *r*_*E*_ and half-maximum activation *k*), natural turnover (with a rate δ_*E*_), and a competition term representing cellular homeostasis due to limited space and resources (with a rate *ε*). The parameter *κ*_*E*_ denotes the rate at which the pathogen causes reduction in the number of active effector T cells able to provide anti-microbial immunity, either via T cell exhaustion, activation-induced cell death, or regulatory T cell intervention. The term *b*. *k* represents the pathogen load at half the maximum inactivation rate of effector T cells, where *b >* 1 is a proportionality constant.

To study the effects of T cell reactivity to pMHC in the context of acute vs. chronic infection, we denoted the spectrum of effector T cells in time by *E*(*t, k*), where *k* = 1/*a*_*k*_ is assumed to be proportional to the reciprocal of T cell reactivity to pMHC consistent with (Standifer *et al*., 2009), *k*_max_ is the maximum value of *k* (corresponding to T cells with the lowest reactivity to pMHC) and *k*_min_ is the minimum value of *k* (corresponding to T cells with highest reactivity to pMHC). Equations (1) and (2) are simulated by discretizing the allowable values of pMHC reactivity within the range defined by 1/*k*_max_ and 1/*k*_min_, wherein the solution is obtained by integrating a high-dimensional system of ordinary differential equations (see Supplementary Text).

### Model parameters and numerical implementation

The two pMHC reactivity dependent parameters, σ_*E*_ and *κ*_*E*_, were defined to be functions of T cell reactivity to pMHC (*a*_*k*_) (**Figure 1B, C**). A bell-shaped function of pMHC reactivity that mimics a log-normal distribution was used to assign values for thymic input σ_*E*_, while T cell exhaustion *κ*_*E*_ was first sampled from an exponential distribution and then sorted in an ascending order (such that higher-affinity T cells are more susceptible to losing their effector functions than their lower-affinity counterparts). This latter claim is supported by the fact that activation-induced cell death (Alexander-Miller *et al*., 1996) and T cell exhaustion (Shakiba *et al*., 2021; Wherry *et al*., 2003) scale with the strength of antigenic stimulation. The use of a uniform distribution for *κ*_*E*_ produced very similar results obtained here (not shown). Table S1 summarizes the meanings of the different parameters and the values used to generate model results. Model parameters were obtained by genetic algorithm fitting to LCMV-Arm and LCMV-Cl13 serum data adapted from (Wherry *et al*., 2003) (**Figure S1A, B**). To simulate the ΔTCR repertoire, the thymus input parameter σ_*E*_ was modulated by an arctan function such that

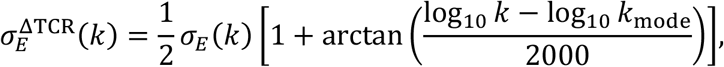

where *k*_mode_ is the mode of the value of the parameter *k* determined by parameter fitting (for more details, see Supplementary Text). The new parameter 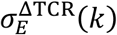 is subsequently renormalized such that the total thymic input across all T cells remains constant, i.e.,

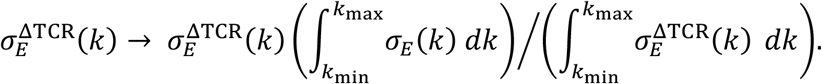

MATLAB was used to simulate the model equations and perform numerical analyses. Steady state and bifurcation analyses were carried out using XPP/AUTO, a freeware available at http://www.math.pitt.edu/~bard/xpp/xpp.html). Genetic algorithm fitting was performed using Compute Canada’s Cedar cluster. MATLAB codes used to produce violin plots are available at https://github.com/bastibe/Violinplot-Matlab. All other MATLAB codes used to perform simulations, parameter fitting, and model analyses can be obtained online (Jamaleddine et al., 2022).

