## Supplemental Figures and Text for "Chronic infection control relies on T cells with lower foreign antigen binding strength generated by N-nucleotide diversity"

**Outline**

Supplementary figures …………………………………………………………………………...……… 2

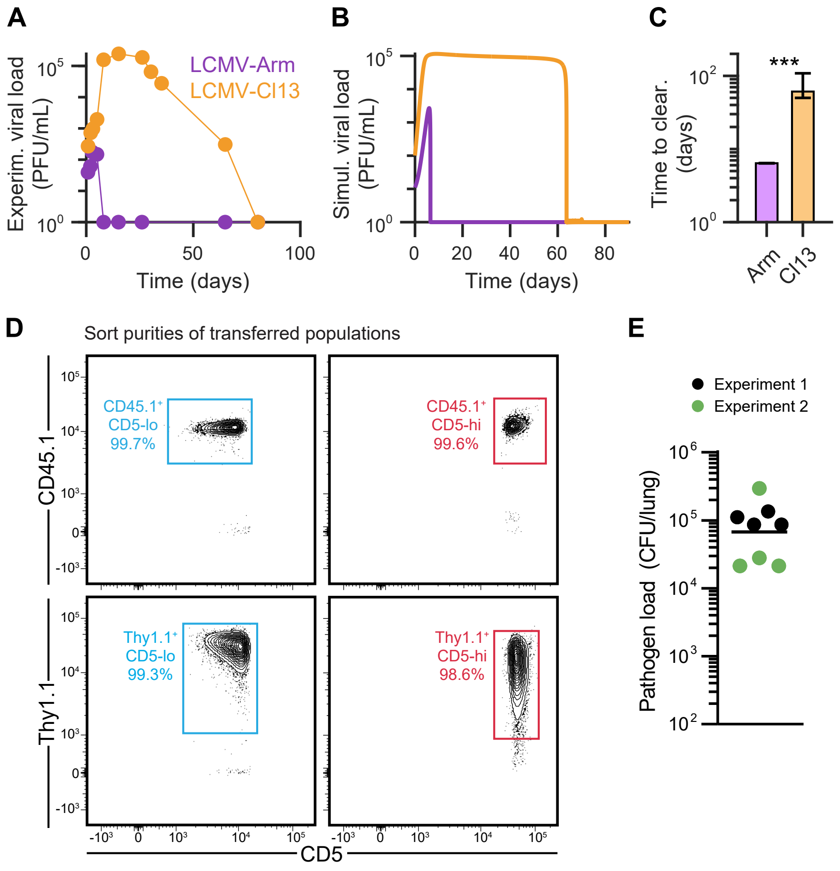

**Figure S1**. Model fitting and experiment set-up details, related to Figure 2. A) Serum viral load data in mice infected with LCMV-Arm or LCMV-Cl13 from (Wherry et al., 2003). B) Model results generated by simulating Eqs. (1) and (2) in the main text using parameters obtained by genetic algorithm fitting of digitized data shown in A (see *Model parameters and fitting*) as listed in Table S1, increasing only the pathogen replication rate parameter. C) Median time to clearance of 100 acute (Arm) or chronic (Cl13) simulated infections (error bars = 95% confidence intervals, $P = 5.24 \times 10^{-8}$ by Wilcoxon rank sum test). D) Flow cytometry panels showing sort purities of transferred $CD45.1^{+}$ or $Thy1.1^{+}$ $CD5^{lo}$ and $CD5^{hi}$ naïve $CD44^{hi}$ $CD62L^{+}$ $CD5^{+}$ T cells into C. neoformans infected mice. E) C. neoformans pathogen loads in the lungs of infected mice from 2 independent experiments (n=8 mice).

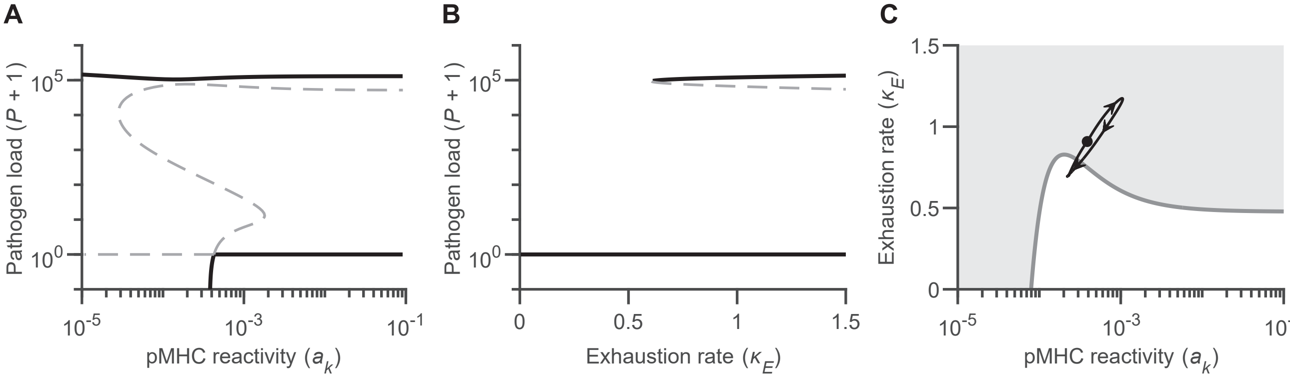

**Figure S2**. Bifurcation analysis of the one-clone model, related to Figure 3. A) Pathogen levels at steady state as a function of pMHC reactivity ($a_k = 1/k$); solid black lines represent branches of attracting (stable) equilibria, while dashed lines represent branches of repelling (unstable) equilibria. The upper and lower levels of pathogen load can coexist (in the form of bistability) in the upper range of pMHC reactivity; which one of these two steady states can be attained depend on the initial conditions of pathogen load and T cell counts. B) Pathogen levels at steady state as a function of the virus-dependent effector T cell depletion, $\kappa_E$, when $a_k = 10^{-3}$; as before, solid black lines represent branches of attracting (stable) equilibria, while dashed lines represen t branches of repelling (unstable) equilibria. C) Two-parameter bifurcation of steady state level of pathogen load with respect to the depletion rate $\kappa_E$ and pMHC reactivity parameter, $a_k$. Gray-shaded region represents the regime of coexistence between the upper and lower levels of pathogen load (i.e., the bistable regime) seen in A and B. Overlayed is the trajectory of the average pMHC reactivity and average depletion rate of the ensemble of T cells of the full system, starting from the filled black circle (arrows indicate direction of motion).

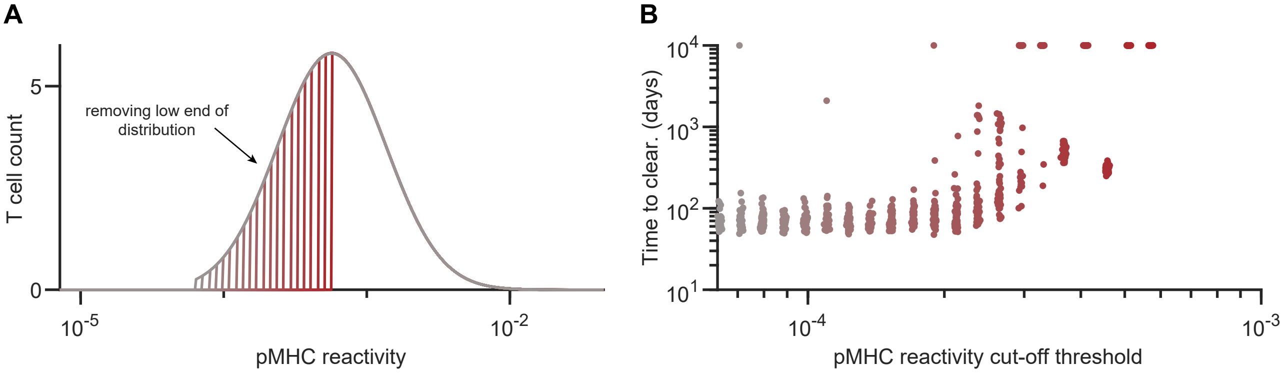

**Figure S3**. Removing low pMHC-affinity T cells, related to Figure 4. A) Distributions of initial T cell count prior to infection as a function of pMHC reactivity obtained by successively removing low-reactivity T cells using different cut-off thresholds from the model's pre-infection repertoire, without keeping the total number of T cells conserved, until only the upper half of the distribution is kept. B) Time to pathogen clearance as a function of the cut-off threshold for T cell reactivity shown in the left panel. For each cut-off threshold, 50 simulation trials were performed as described in Figure 4. Notice that the acute cluster has disappeared compared to Figure 4B.

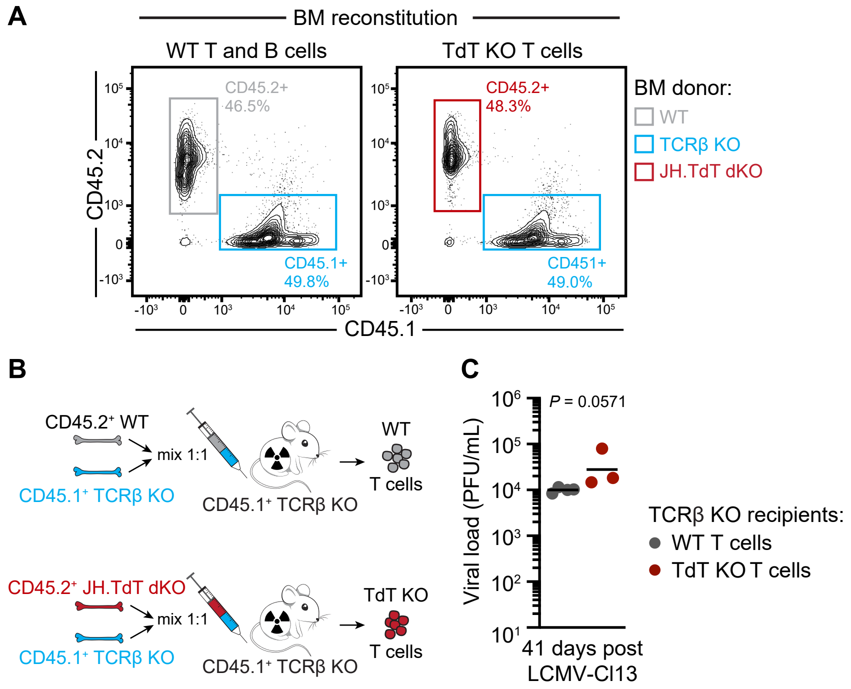

**Figure S4**. Generating bone marrow chimeric mice lacking TdT in T cells only, related to Figure 5. A) Representative flow cytometry plots showing the percent of bone marrow cells from each set of donor mice, namely CD45.1$^{+}$ cells from TCRβ KO mice and CD45.2$^{+}$ cells from either WT or JH x TdT double KO mice. B) Modified experimental approach from Figure 5A by using TCRβ KO mice as irradiated bone marrow recipients. C) LCMV-Cl13 viral loads in the serum of recipient TCRβ$^{-/-}$ mice reconstituted with WT or TdT KO T cells in at day 41 post infection. P value indicated was computed using two-tailed Wilcoxon rank sum test on geometric means.

### Model parameters and fitting

**Table S1.** Parameter values used in model simulations. Values are determined by fitting model simulations to serum data in acute vs. chronic LCMV infection (Wherry *et al.*, 2003) using a genetic algorithm.

| Parameter | Description | Value(s) |
|---|---|---|
| $r_P$ | Maximum pathogen replication rate | 1.31 day$^{-1}$ (acute), 1.91 day$^{-1}$ (chronic) |
| $P_{\max}$ | Pathogen carrying capacity | $1.82\times10^{5}$ PFU mL$^{-1}$ |
| $\kappa_P$ | Maximum pathogen removal rate per T cell | $1.91\times10^{-2}$ PFU mL$^{-1}$ cell$^{-1}$ day$^{-1}$ |
| $k$ | Viral load for half-maximum activation of T cells | Range |
| $k_{\min}$ | Minimum value of $k$ corresponding to highest T-cell avidity | 22.1 pfu mL$^{-1}$ |
| $k_{\max}$ | Maximum value of $k$ corresponding to lowest T-cell avidity | $1.39\times10^{5}$ pfu mL$^{-1}$ |
| $a_k$ | pMHC reactivity parameter (equal to $1/k$) | Range |
| $a$ | Scaling factor of half-maximum constant in viral clearance | $3.1\times10^{-3}$ |
| $\sigma_E$ | Thymic input, source of naïve effector T cells | 122.4 cells day$^{-1}$ (total) |
| | Assigned as a theoretical log-normal distribution $f(k;\ \mu,\sigma)$ | $\mu$=8.24, $\sigma$=0.88 (WT) |
| | | $\mu$=7.38, $\sigma$=0.36 (TdT KO) |
| $r_E$ | Proliferation rate of effector T cells | 3.79 day$^{-1}$ |
| $\delta_E$ | Natural turnover rate of effector T cells | 0.17 day$^{-1}$ |
| $\kappa_E$ | Virus-dependent depletion rate of effector T cells (e.g., exhaustion) | |
| | Sampled from shifted exponential distribution $g(k;\mu)+\kappa_{E,\min}$ and sorted (highest $\alpha_k$ corresponds to highest $\kappa_E$ value) | $\mu$=1.06, $\kappa_{E,\min}$=0.2 day$^{-1}$ |
| $b$ | Scaling factor of half-maximum constant in T cell depletion | 35.8 |
| $\varepsilon$ | Competition rate due to homeostasis | $6.91\times10^{-7}$ cell$^{-1}$ day$^{-1}$ |
| $N$ | Number of T-cell clones obtained from the discretization of system of Eqs. (1)-(2) (see *Numerical simulation* section) | 500 |

In order to generate parameter values for model simulations, a genetic algorithm is used. The fitness function of the genetic algorithm is primarily based on minimizing of the sum of squared errors (i.e., minimizing $\sum_i \big(\boldsymbol{Y}_i - F(t;\boldsymbol{P})\big)^2$ where $\boldsymbol{Y}_i$ is the serum time series data in (Wherry *et al.*, 2003), $F(t,\boldsymbol{P})$ is the fit output as a function of time with input parameter vector $\boldsymbol{P}$) is the between the data and model predictions. The results are then refined to ensure the final selected parameter values satisfy the following conditions:

- The pathogen replication rate for chronic LCMV (LCMV-Cl13) is larger than that for acute LCMV (LCMV-Arm), as suggested by (Bergthaler et al., 2010; Sullivan et al., 2011)
- The rate of change of pathogen is initially positive, i.e., $\mathrm{d}P/dt > 0$ at $V = 0$.
- The steady state of the pathogen-free equilibrium is stable (see "*Stability analysis"* section below)
- An upper bound for the standard deviation the function describing the $\sigma_E$ distribution, $f(k;\ \mu,\sigma)$, is applied to ensure that $f$ is negligibly small for non-physiologically small values of $k$ (i.e., for pMHC reactivities that are too high).

### Numerical simulation

To simulate the system of integro-differential equations, we discretize the continuous T cell population $E(t,k)$, into individual clones of T cells $E_i$ with pMHC reactivity $a_{k,i} = 1/k_i$, where $i = 1,2,\ldots,N$ and $N$ is the number of T cell clonotypes. To approximate an pMHC-reactivity continuum, we choose a large value for $N$. The resulting model becomes a high-dimensional system of ordinary differential equations with $N+1$ variables, whose equations can be expressed as

$$\frac{\mathrm{d}P}{\mathrm{d}t} = r_P\, P\left(1 - \frac{P}{P_{\max}}\right) - \kappa_P \sum_{i=1}^{N} E_i \frac{P}{P + a\, k_i}$$

$$\frac{\mathrm{d}E_i}{\mathrm{d}t} = \sigma_{E_i} + r_E\, E_i\, \frac{P}{P + k_i} - \delta_E\, E_i - \kappa_E\, E_i\, \frac{P}{P + b\, k_i} - \varepsilon\, E_i \sum_{j=1}^{N} E_j$$

### Stability analysis of the full model

One can show that $(P, \boldsymbol{E}) = (0, \boldsymbol{E}^*)$, where $\boldsymbol{E}^* = (E_1^*, E_1^*, \ldots, E_N^*)$ is the level of each effector T cell clone in the pathogen-free equilibrium, is always a steady state solution of the full, discretized system. We denote this steady state by $\boldsymbol{S}_0$. In this section, we will determine the condition needed to ensure that $\boldsymbol{S}_0$ is stable. This condition is then imposed on the genetic algorithm when evaluating parameter fitness.

The Jacobian matrix of the full system evaluated at the steady state $\boldsymbol{S}_0 = (0, \boldsymbol{E}^*)$ can be written as:

$$\mathbf{J}_{\boldsymbol{S}_0} = \begin{pmatrix} r_P - \dfrac{\kappa_P}{a}\displaystyle\sum_{n=1}^{N}\frac{E_n^*}{k_n} & 0 & 0 & \cdots & 0 \\ \dfrac{E_1^*}{k_1}\left(r_E - \dfrac{\kappa_E}{b}\right) & -\delta_E - \varepsilon\displaystyle\sum_{n=1}^{N} E_n^* - \varepsilon E_1^* & -\varepsilon E_1^* & \cdots & -\varepsilon E_1^* \\ \dfrac{E_2^*}{k_2}\left(r_E - \dfrac{\kappa_E}{b}\right) & -\varepsilon E_2^* & -\delta_E - \varepsilon\displaystyle\sum_{n=1}^{N} E_n^* - \varepsilon E_2^* & \cdots & -\varepsilon E_2^* \\ \vdots & \vdots & \vdots & \ddots & \vdots \\ \dfrac{E_N^*}{k_N}\left(r_E - \dfrac{\kappa_E}{b}\right) & -\varepsilon E_N^* & -\varepsilon E_N^* & \cdots & -\delta_E - \varepsilon\displaystyle\sum_{n=1}^{N} E_n^* - \varepsilon E_N^* \end{pmatrix}$$

It can be shown that the eigenvalues of the system can be written as

$$\lambda_0 = r_P - \frac{\kappa_P}{a}\sum_{n=1}^{N}\frac{E_n^*}{k_n}, \qquad \lambda_1 = -\delta_E - 2\varepsilon\sum_{n=1}^{N} E_n^*, \qquad \lambda_i = -\delta_E - \varepsilon\sum_{n=1}^{N} E_n^*, \qquad i = \{2,3,\ldots,N\}.$$

Therefore, $\lambda_0 < 0$ is a necessary and sufficient condition to ensure local stability of $\boldsymbol{S}_0$, as all other eigenvalues are always negative for positive parameter values and T-cell levels. This condition may be rewritten as

$$\sum_{n=1}^{N} \frac{E_n^*}{k_n} > \frac{a r_P}{\kappa_P}.$$

The individual values of $E_n^*$ may be computed by simulating the full model in the absence of pathogen (i.e., for $P = 0$); this condition can then be verified during the parameter fitness evaluation process of the genetic algorithm to reject parameter selections that do not ensure the stability of $\boldsymbol{S}_0$.

The existence and stability of other steady states in the model depends on different parameter values. To assess them, we employ bifurcation analysis of a simplified single-clone ($N = 1$), 2-dimensional model.

**Analysis of the reduced one-clone, 2D model**

To better understand the underlying dynamics of the full continuum model and the distinct time scales between outcomes of an acute and chronic infection, we turn out attention to a simplified, single-clone version of the model (with $N = 1$), given by

$$\frac{\mathrm{d}P}{\mathrm{d}t} = r_P\, P\left(1 - \frac{P}{P_{\max}}\right) - \kappa_P\, E\,\frac{P}{P + a\,k}$$
$$\frac{\mathrm{d}E}{\mathrm{d}t} = \sigma_E + r_E\, E\,\frac{P}{P + k} - \delta_E\, E - \kappa_E\, E\,\frac{P}{P + b\,k} - \varepsilon\, E^2.$$

Model parameters are assigned the same values as those provided in Table S1, except for $\sigma_E$, $\kappa_E$ and $k$. In this case, $\sigma_E$ is set to be 122.4 cells/day, $\kappa_E$ to be 1.06 day$^{-1}$, and $k$ to be a bifurcation parameter spanning the entire range $[k_{min}, k_{max}]$.

Figure S2A plots the steady-state levels of the pathogen load, $P$, on a log-scale, as a function of the pMHC reactivity ($\alpha_k$). The pathogen load is shifted by 1, to show the behaviour at $P = 0$, the "undetectable" level of pathogen load, on a log-scale (i.e., $10^0$ corresponds to $P = 0$). The figure shows that pathogen load can attain two different values at steady state (solid lines): an elevated level and a low level, both acting as attractors (as opposed to repellers shown as dashed lines) that can co-exist at high $\alpha_k$. This coexistence of the two attractors is a hallmark of bistability, which highlights the dependence of the system on the initial level of $P$ to determine which attractor will be eventually approached over time. For the $P = 0$ steady state to be an attractor (stable), the condition $\alpha_k > 4.27 \times 10^{-4}$ must be satisfied.[a]

Choosing $a_k = 1/k = 10^{-3}$ such that $P = 0$ is stable, we next show the behaviour of the system with respect to the pathogen-dependent effector T-cell exhaustion rate, $\kappa_E$ (Fig. S2B).

[a] Note that units are omitted from the discussion of the pMHC-reactivity measure, since $\alpha_k$ does not represent a direct value for the affinity of the T cell receptor but rather acts as an indicator for it.

Pathogen loads can be either elevated or zero due to the presence of bistability for $\kappa_E > 0.6$ day$^{-1}$. Below this critical value of $\kappa_E$, only the lower steady state exists as the system's global attractor.[b]

To understand the effects of such dynamics of the single-clone model on the full continuum model, we compute the evolution of the weighted average of pMHC reactivity and that of the exhaustion rate of dominant effector T cells throughout the chronic immune response and overlay this trajectory on the 2-parameter bifurcation diagram of viral load with respect to $\kappa_E$ and $\alpha_k$ of the single clone model (Fig. S2C). The gray-shaded region represents the bistable region in Fig. S2B. The starting values for the average depletion rate and avidities falls within the bistable region; acute vs. chronic outcomes are set apart by whether the system evolves immediately toward the $P = 0$ steady state (acute), or if it first goes to the upper steady state and can only come back down once the full system moves out of the gray region. This latter process will take much longer, hence the distinct time scale separations observed in the time to clearance of the full model that clearly defines acute vs. chronic pathogen replication durations.

[b] Within the physiological range of the system, $P \geq 0$ and $E \geq 0$.
